# Evaluating potential drug targets through human loss-of-function genetic variation

**DOI:** 10.1101/530881

**Authors:** Eric Vallabh Minikel, Konrad J Karczewski, Hilary C Martin, Beryl B Cummings, Nicola Whiffin, Daniel Rhodes, Jessica Alföldi, Richard C Trembath, David A van Heel, Mark J Daly, Genome Aggregation Database Production Team, Genome Aggregation Database Consortium, Stuart L Schreiber, Daniel G MacArthur

## Abstract

Naturally occurring human genetic variants predicted to cause loss of function of protein-coding genes provide an *in vivo* model of human gene inactivation that complements cell and model organism knockout studies. Here we investigate the application of human loss-of-function variants to assess genes as candidate drug targets, with three key findings. First, even essential genes, where loss-of-function variants are not tolerated, can be highly successful as targets of inhibitory drugs. Second, in most genes, loss-of-function variants are sufficiently rare that genotype-based ascertainment of homozygous or compound heterozygous “knockout” humans will await sample sizes ~1,000 times those available at present. Third, automated variant annotation and filtering are powerful, but manual curation remains critical for removing artifacts and making biological inferences, and is a prerequisite for recall-by-genotype efforts. Our results provide a roadmap for human “knockout” studies and should guide interpretation of loss-of-function variants in drug development.

## MAIN TEXT

Human genetics is an increasingly critical source of evidence guiding the selection of new targets for drug discovery^1^. Most drug candidates that enter clinical trials eventually fail for lack of efficacy^2^, and while *in vitro*, cell culture, and animal model systems can provide preclinical evidence that the compound engages its target, too often the target itself is not causally related to human disease^1^. Candidates targeting genes with human genetic evidence for disease causality are more likely to become approved drugs^3,4^, and identification of humans with loss-of-function (LoF) variants, particularly two-hit (homozygous or compound heterozygous) genotypes, has, for several genes, correctly predicted the safety and phenotypic effect of pharmacologically inhibiting the drug’s target^5^. While these examples demonstrate the value of human genetics in drug development, important questions remain regarding strategies for identifying individuals with LoF variants in a gene of interest, interpretation of the frequency — or lack — of such individuals, and whether it is wise to pharmacologically target a gene in which LoF variants are associated with a deleterious phenotype.

Public databases of human genetic variation have provided catalogs of predicted loss-of-function (pLoF) variants — nonsense, essential splice site, and frameshift variants expected to result in a non-functional allele — in humans. Such databases present a new opportunity to study the effects of pLoF variation in genes of interest and to identify individuals with pLoF genotypes in order to understand gene function or disease biology, or to assess potential for therapeutic targeting. While many variants initially annotated as pLoF do not, in fact, cause a loss of function^6^, rigorous automated filtering can remove common error modes^7^. True LoF variants are generally rare, and show important differences between outbred, bottlenecked^8^, and consanguineous^9^ populations^6,10^. Counting the number of distinct pLoF variants in each gene in a population sample allows quantification of gene essentiality in humans through a metric known as *constraint*^10–13^. Specifically, the rate at which *de novo* pLoF mutations arise in each gene is predicted based on DNA mutation rates^10,12^, and the ratio of the count of pLoF variants observed in a database to the number expected based on mutation rates — obs/exp or simply constraint score — measures how strongly purifying natural selection has removed such variants from the population. Annotation of pLoF variants remains imperfect, and continued improvements are being made^14^, but the fact that constraint usefully measures gene essentiality is demonstrated by agreement with cell culture and mouse knockout experiments^7^, by overlap with human disease genes^7,10^ and genes depleted for structural variation^15^, and by the power of constraint to enrich for deleterious variants in neurodevelopmental disorders^7,16^.

Building on these insights, here we leverage pLoF variation in the Genome Aggregation Database (gnomAD)^7^ v2 dataset of 141,456 individuals to answer open questions in the interpretation of human pLoF variation in disease biology and drug development.

### Constraint in human drug targets

We compared constraint in the targets of approved drugs extracted from DrugBank^17^ (*N*=383) versus all protein-coding genes *(N*=17,604). Drug targets were, on average, just slightly more constrained than all genes (mean 44% vs. 52%, *P*=0.00028), but the two gene sets had a qualitatively similar distribution of scores, ranging from intensely constrained (0% obs/exp) to not at all constrained (≥100% obs/exp; Fig. 1a). Constraint scores showed clear divergence between categories of genes (Extended Data Table 1) expected to be more or less tolerant of inactivation (Fig. 1b), as previously reported^7,10^, validating the usefulness of constraint as a measure of gene essentiality. Nonetheless, when drug targets were stratified by drug modality, indication, or effect, no statistically significant differences between subsets of drug targets were observed.

**Figure 1.**
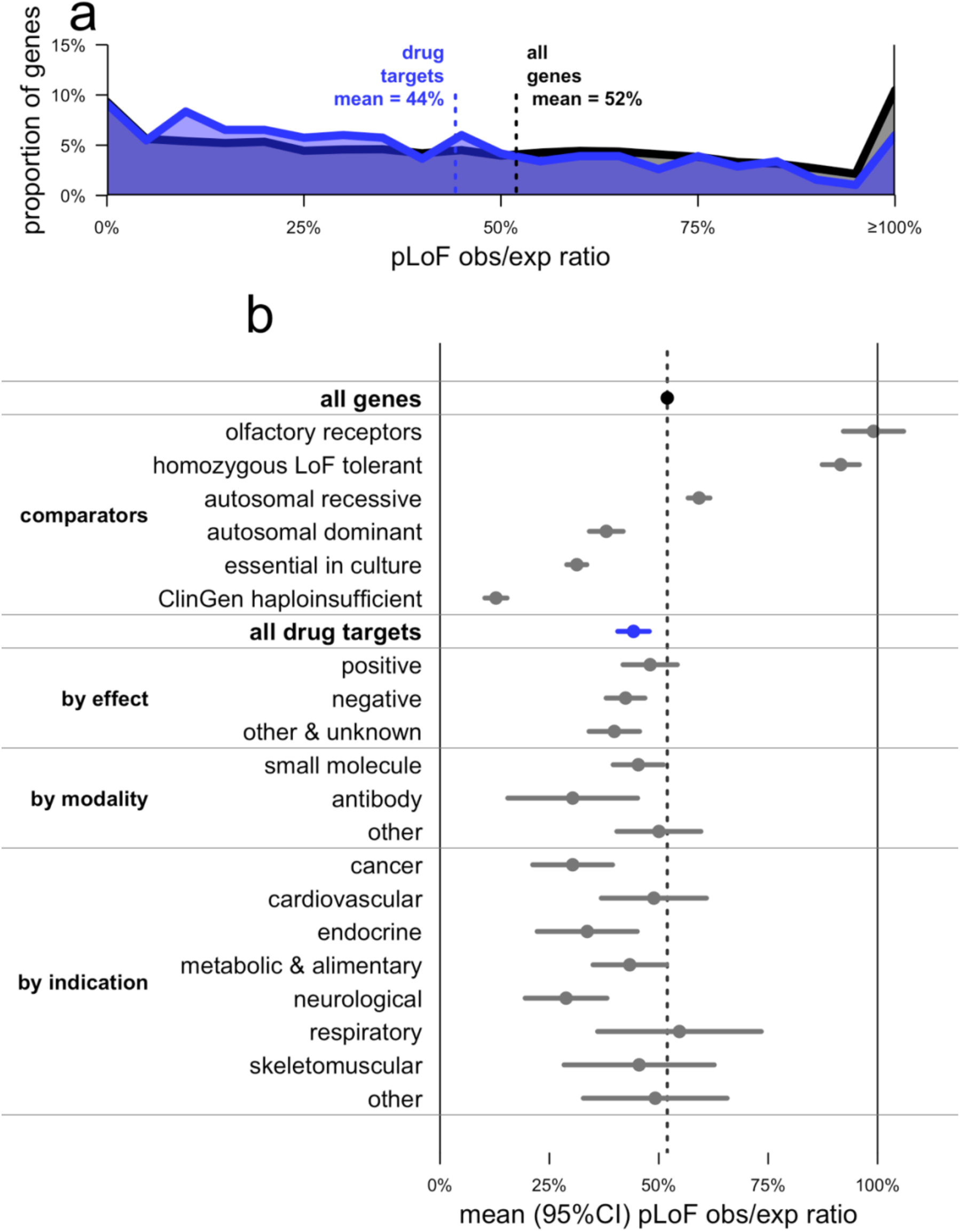
pLoF constraint in drug targets. **a)** Histogram of pLoF obs/exp for all genes (black) versus drug targets (blue). **b)** Forest plot of means (dots) and 95% confidence intervals indicating our certainty about the mean (line segments), for pLoF obs/exp ratio in the indicated gene sets. See Online Methods for data sources. For drug effect, ‘positive’ indicates agonist, activator, inducer, etc., while negative indicates antagonist, inhibitor, suppressor, etc.

The slightly but significantly lower obs/exp value among drug targets may superficially appear to provide evidence that constrained genes make superior drug targets. Stratification of drug targets by protein family, human disease association, and tissue expression, however, argues against this interpretation. Drug targets are strongly enriched for a few canonically “druggable” protein families, for genes known to be involved in human disease, and for genes with tissue-restricted expression; each of these properties is in turn correlated with either significantly stronger or weaker constraint (Extended Data Fig. 1). Although controlling for these correlations does not abolish the trend of stronger constraint among drug targets, the correlation of so many observed variables with a gene’s status as drug target argues that many unobserved variables likely also confound interpretation of the lower mean obs/exp value among drug targets.

The overall constraint distribution of drug targets (Fig. 1a) also argues against the view that a gene in which LoF is associated with a deleterious phenotype cannot be successfully targeted. Indeed, 19% of drug targets (*N*=73), including 52 targets of inhibitors, antagonists or other “negative” drugs, have obs/exp values lower than the average (12.8%) for genes known to cause severe diseases of haploinsufficiency^18^ (ClinGen Level 3). To determine whether this finding could be explained by particular class or subset of drugs, we examined constraint in several well-known example drug targets (Table 1). Some heavily constrained genes are targets of cytotoxic chemotherapy agents such as topoisomerase inhibitors or cytoskeleton disruptors, a set of drugs intuitively expected to target essential genes. However, genes with near-complete selection against pLoF variants also include *HMGCR* and *PTGS2*, the targets of highly successful, chronically used inhibitors — statins and aspirin.

**Table 1.** Spectrum of tolerance to genetic inactivation among human drug targets. Example targets are arranged from most intolerant of inactivation (top) to most tolerant (bottom).

| drug class | example | gene | obs/exp pLoF |
| --- | --- | --- | --- |
| topoisomerase I inhibitors | irinotecan | *TOP1* | 0% (0/50.5) |
| M1-selective antimuscarinics | pirenzepine | *CHRM1* | 0% (0/14.1) |
| cytoskeleton disruptors | paclitaxel | *TUBB* | 6% (1/16.4) |
| non-steroidal anti-inflammatory drugs (NSAIDs) | aspirin | *PTGS2* | 10% (3/29.7) |
| statins | atorvastatin | *HMGCR* | 13% (6/46.3) |
| phosphodiesterase 5 inhibitors | sildenafil | *PDE5A* | 33% (16/47.8) |
| antifolates | methotrexate | *DHFR* | 38% (4/10.5) |
| proton pump inhibitors | omeprazole | *ATP4A* | 52% (25/47.9) |
| antiplatelets | clopidogrel | *P2RY12* | 66% (5/7.6) |
| H1 antihistamines | cetirizine | *HRH1* | 76% (11/14.5) |
| angiotensin converting enzyme (ACE) inhibitors | benazepril | *ACE* | 87% (62/71.3) |
| PCSK9 antibodies | alirocumab | *PCSK9* | 98% (26/26.5) |

These human *in vivo* data further the evidence from other species and models that essential genes can be good drug targets. Homozygous knockout of *Hmgcr* and *Ptgs2* are lethal in mice^19–21^. Drug targets exhibit higher inter-species conservation than other genes^22^. Targets of negative drugs include 14 genes with lethal heterozygous knockout mouse phenotypes reported^23^ and 6 reported as essential in human cell culture^24^.

### Prospects for finding “knockout” individuals

While constraint alone is not adequate to nominate or exclude drug targets, the study of individuals with single hit (heterozygous) or two-hit (“knockout”) LoF genotypes in a gene of interest can be highly informative about the biological effect of engaging that target^5^. To assess prospects for ascertaining “knockout” individuals, we computed the cumulative allele frequency (CAF) of pLoF variants in each gene (Online Methods), and then used this to estimate the expected frequency of two-hit individuals under different population structures (Fig. 2) absent natural selection.

**Figure 2.**
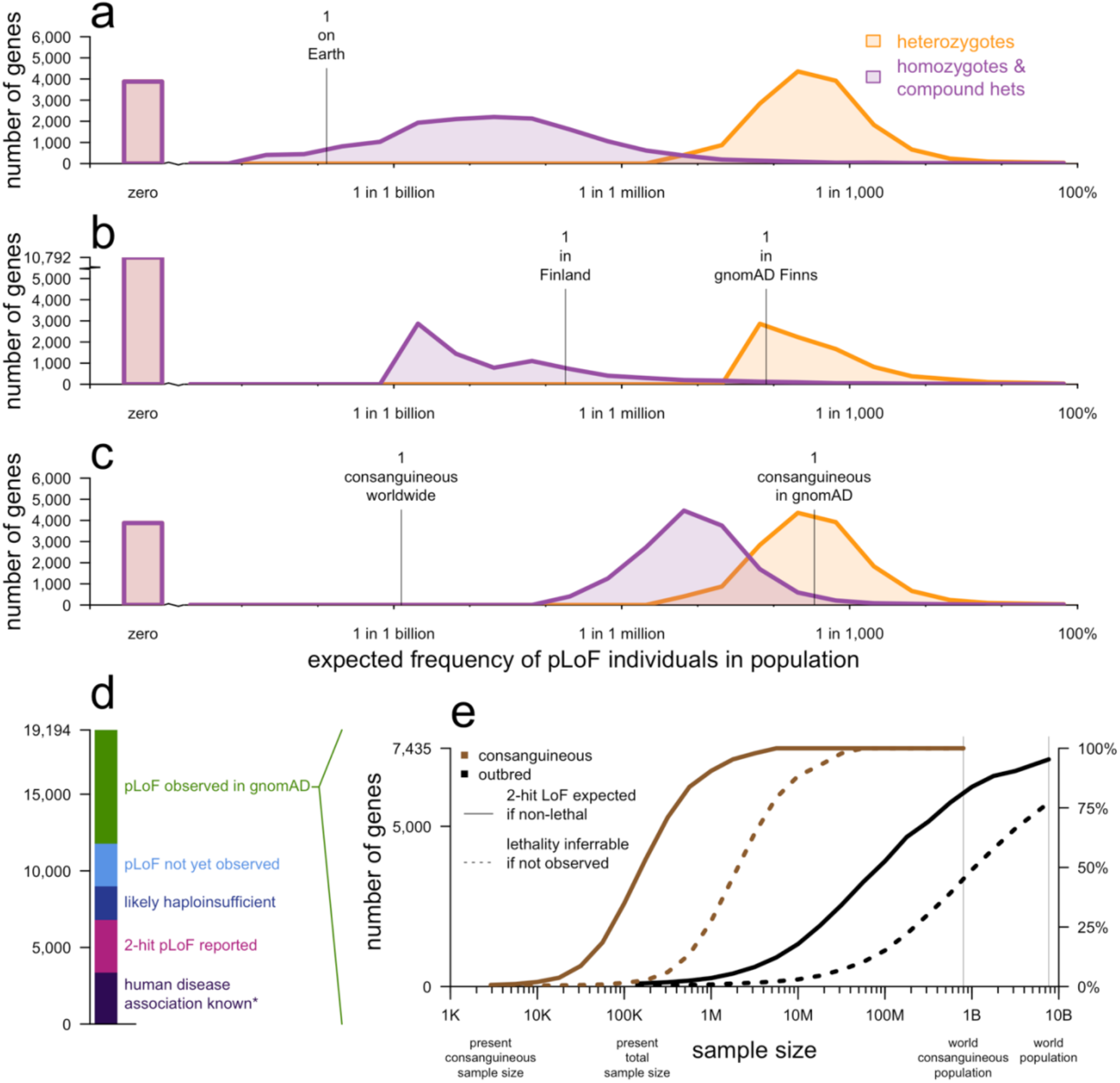
Prospects for discovery of human “knockouts”. Each panel **a-c** shows a histogram where the y axis is number of genes and the x axis shows the theoretically expected population frequency of single hit heterozygotes (orange), versus two-hit homozygotes and compound heterozygotes (purple). Zero indicates the number of genes where no pLoF variants have been observed. **a)** Outbred populations, under random mating — heterozygotes have frequency 2p(1-p) and two-hit individuals have frequency p^2^. The value of p is taken from all gnomAD exomes. **b)** Finnish individuals, an example of a bottlenecked population. Single and two-hit frequencies are again 2p(1-p) and p^2^ but p is based on Finnish exomes only. **c)** Consanguineous individuals with a = 5.8% of their genome autozygous (both chromosomes inherited from the same recent ancestor); heterozygote frequency is 2p(1-p) and two-hit frequency is (1-a)p^2^ + ap. See Online Methods for details. **d)** Current status of pLoF or disease association discovery for all protein-coding genes. **e)** For genes with pLoF observed in gnomAD (top category from d), projected sample sizes required for discovery of two-hit individuals (solid lines) and for statistical inference that a two-hit genotype is lethal if no such individuals are observed (dashed lines), for consanguineous and outbred individuals.

Whereas gnomAD is now large enough to include at least one pLoF heterozygote for the majority (15,317/19,194; 79.8%) of genes, ascertainment of total “knockout” individuals in outbred populations will require 1,000-fold larger sample sizes for most genes: the median gene has an expected two-hit frequency of just 6 per billion (Fig. 2a). Even if every human on Earth were sequenced, there are 4,728 genes (25%) for which identification of even one two-hit individual would not be expected in outbred populations. Intuitively, because today’s gnomAD sample size is larger than the square root of the world population, variants so far seen in zero or only a few heterozygous individuals are not likely to ever be seen in a homozygous state in outbred populations, except where variants prove common in populations not yet well-sampled by gnomAD.

Because population bottlenecks can result in very rare variants present in a founder rising to an unusually high frequency, we also considered the utility of bottlenecked populations for knockout discovery, using Finnish individuals in gnomAD as an example^8^. Although this population structure can enable well-powered association studies for the small fraction of genes in which pLoF variants drifted to high frequency due to the bottleneck, overall, identification of two-hit pLoF individuals for a pre-specified gene of interest appears equally or more difficult in Finns than in outbred populations (Fig. 2b and Extended Data Fig. 2), because rare variants not present in a founder have been effectively removed from the population.

Finally, we considered consanguineous individuals, where parental relatedness greatly increases the frequency of homozygous pLoF genotypes. The *N*=2,912 individuals in the East London Genes & Health (ELGH) cohort^25^ who report having parents who are second cousins or closer have on average 5.8% of their genomes autozygous. Here, the expected frequency of two-hit individuals is many times higher than in outbred populations, at 5 per million for the median gene (Fig. 2c).

These projections allow us to draft a roadmap for discovery of human knockouts (Fig. 2d-e). Of 19,194 genes, 3,367 (18%) already have a human disease association annotated in OMIM, although we note that the discovery of LoF individuals in population databases will still be valuable for assessing penetrance and for identifying LoF syndromes associated with known GoF genes. There are 3,421 genes (18%) without known human disease association that have two-hit pLoF genotypes reported in gnomAD^7^, ELGH^26^, PROMIS^27^, DeCODE^28^, or UK BioBank^29^, suggesting this genotype may be tolerated. An additional 2,190 genes (11%) can be inferred likely intolerant of heterozygous inactivation (pLI > 0.9) in gnomAD, and would be expected to be enriched for genes with severe heterozygous and lethal homozygous LoF phenotypes. Another 2,781 genes (14%) have no pLoF variants yet observed in gnomAD, but our sample size is not yet large enough to robustly infer LoF intolerance. For these genes, observation of outbred two-hit individuals is not expected, and we cannot yet assess the feasibility of identifying consanguineous two-hit individuals because we lack an estimate of pLoF allele frequency.

This leaves 7,435 genes (39%) for which one or more pLoFs are observed in gnomAD, but strong LoF intolerance cannot be inferred, nor have two-hit genotypes been observed, nor is a human disease phenotype known. We projected the sample sizes required to identify “knockout” individuals for these genes (Fig. 2e). In outbred populations, current sample size would need to be increased by approximately 1,000-fold before ascertainment of a single two-hit LoF individual would be expected for the typical gene. In contrast, a ~10-to 100-fold increase from current consanguineous sample size, meaning hundreds of thousands of individuals in absolute terms, would identify at least one two-hit LoF individual for the typical gene.

These calculations are based on variants annotated as predicted LoF in gnomAD. Structural and non-coding variation resulting in a loss of function may be missed in exomes, and missense variants resulting in a loss of function cannot be rigorously annotated, leading to underestimation of cumulative LoF allele frequency. Overall, however, our calculations likely represent an upper bound on the total frequency of two-hit individuals in the population. The variants included in this analysis are filtered but have not been manually curated or functionally validated, so some will ultimately prove not to be true LoF. These false positives tend to be more common and will have disproportionately contributed to the cumulative LoF allele frequency. More importantly, for some genes, complete knockout will not be tolerated. When only one or a few two-hit individuals are expected in a dataset, the absence of any such individuals can be due to either early lethality, a severe clinical phenotype incompatible with inclusion in gnomAD, or simply chance. Thus, the ability to infer lethality of this genotype based on statistical evidence will lag behind the identification of two-hit individuals where they do exist (Fig. 2e). For some genes, inference of lethality will always remain impossible in outbred populations, though it may be feasible in consanguineous individuals.

### Curation of pLoF variants

Where pLoF variants can be identified, they are a valuable resource for assessing the impact of lifelong reduction in gene dosage. To highlight the challenges and opportunities of identifying such variants, we manually curated gnomAD data and the scientific literature for six genes associated with gain-of-function (GoF) neurodegenerative diseases, for which inhibitors or suppressors are under development^30–35^: *HTT* (Huntington disease), *MAPT* (tauopathies), *PRNP* (prion disease), *SOD1* (amyotrophic lateral sclerosis), and *LRRK2* and *SNCA* (Parkinson disease). The results (Table 2 and Fig. 3) illustrate four points about pLoF variant curation.

**Table 2.** Curation of pLoF variation in six neurodegenerative disease genes. Shown are the coding sequence length (base pairs, bp), constraint value (pLoF obs/exp) after filtering and curation, cumulative allele frequency before and after filtering and manual curation, estimated frequency of true pLoF heterozygotes in the population, and genetic prevalence (population frequency including pre-symptomatic individuals) of the gain-of-function (GoF) disease associated with the gene. Genetic prevalence calculations are described in Extended Data Table 2, and variant curation details are provided in Supplementary Table 1, except for LRRK2 which is described in detail in Whiffin et al^49^. *Constitutive brain-expressed exons only. **PRNP codons 1-144, see Fig. 3c for rationale.

| gene | length (bp) | pLoF obs/exp | cumulative pLoF allele frequency |  | pLoF heterozygote frequency | GoF disease genetic prevalence |
| --- | --- | --- | --- | --- | --- | --- |
|  |  |  | before filtering & curation | after filtering & curation |  |  |
| *HTT* | 9,426 | 8.2% | 6.2% | 0.013% | 1 in 3,800 | 1 in 2,400-4,400 |
| *LRRK2* | 7,581 | 41% | 0.23% | 0.09% | 1 in 500 | 1 in 3,300 |
| *MAPT* | 2,328 | 0%* | 14% | 0% | — | 1 in 5,000 – 31,000 |
| *PRNP* | 759 | 99%** | 0.0035% | 0.0021% | 1 in 18,000 | 1 in 50,000 |
| *SNCA* | 420 | 0% | 0.0012% | 0% | — | 1 in 360,000 |
| *SOD1* | 462 | 18% | 0.0060% | 0.0038% | 1 in 26,000 | 1 in 27,000-83,000 |

**Figure 3.**
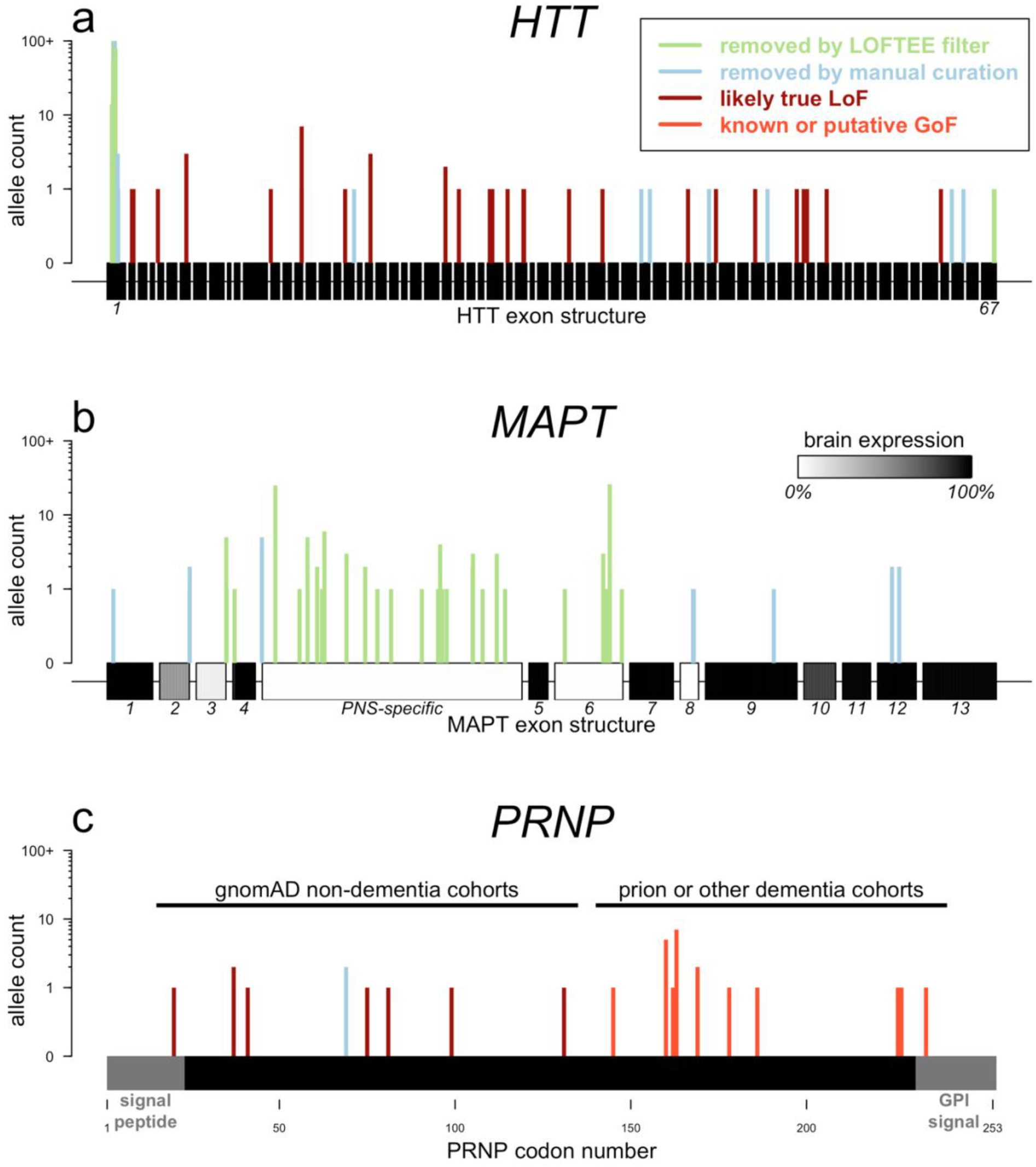
Insights from non-random positional distributions of pLoF variants. **a)** HTT, **b)** MAPT, with exon numbering and annotation from Andreadis^51^ and brain expression data from GTEx^40^, and **c)** PRNP, a single protein-coding exon with domains removed by post-translational modification in gray, showing previously reported variants^41^ as well as those newly identified in gnomAD and in the literature^52,53^. See text for interpretation, Extended Data Table 3 for details of PRNP variants, and Supplementary Table 1 for detailed variant curation results.

First, other things being equal, genes with longer coding sequences offer more opportunity for LoF variants to arise, and so tend to have a higher cumulative frequency of LoF variants, unless they are heavily constrained. Ascertainment of LoF individuals is thus harder for shorter and/or more constrained genes, even though these may be good targets (Table 1).

Second, many variants annotated as pLoF are false positives^6^, and these are enriched for higher allele frequencies, so that both filtering and curation have an outsized impact on the cumulative allele frequency of LoF. Studies of human pLoF variants lacking stringent curation can therefore easily dilute results with false pLoF carriers.

Third, after careful curation, cumulative LoF allele frequency is sometimes sufficiently high to place certain bounds on what heterozygote phenotype might exist. For example, GoF mutations causing genetic prion disease have a ~1 in 50,000 genetic prevalence^36^ and have been known for three decades, with thousands of cases identified, making it unlikely that a comparably severe and penetrant haploinsufficiency syndrome associated with *PRNP* would have gone unnoticed to the present day despite being more than twice as common (~1 in 18,000). Similar arguments can be made for *HTT*, *LRRK2*, and *SOD1*. Of course, this does not rule out the possibility that heterozygous loss-of-function in these genes could be associated with less severe or less penetrant phenotypes.

Finally, careful inspection of the distributions of pLoF variants can reveal important error modes or disease biology. *HTT*, *MAPT*, and *PRNP* each have different non-random positional distributions of pLoF variants (Fig. 3). High-frequency *HTT* pLoF variants cluster in the polyglutamine/polyproline repeat region of exon 1 and appear to be alignment artifacts (Fig. 3a). True *HTT* LoF variants are rare and the gene is highly constrained, which might suggest some fitness effect in a heterozygous state in addition to the known severe homozygous phenotype^37,38^, although the frequency of LoF carriers still argues against a penetrant syndromic illness, consistent with the lack of phenotype reported in heterozygotes identified to date^38,39^. High-frequency *MAPT* pLoF variants cluster in exons not expressed in the brain in GTEx data^14,40^, and all remaining pLoFs appear to be alignment or annotation errors (Fig 3b). No true LoFs are observed in *MAPT*, although our sample size is insufficient to prove that *MAPT* LoF is not tolerated — among constitutive brain-expressed exons, we expect 12.6 LoFs and observe 0, giving a 95% confidence interval upper bound of 23.7% for obs/exp. *PRNP* truncating variants in gnomAD cluster in the N terminus; the sole C-terminal truncating variant in gnomAD is a dementia case (Extended Data Table 2), consistent with variants at codon ≥145 causing a pathogenic gain-of-function through change in localization (Fig. 3c). Within codon 1-144, *PRNP* is unconstrained, and no neurological phenotype has been identified in individuals with truncating variants to date, consistent with the hypothesis that N-terminal truncating variants are true LoF and are tolerated in a heterozygous state^41^.

## Discussion

The study of gene inactivation through human genetic databases can illuminate human biology and guide drug target selection, complementing mouse knockout studies^42^, but analysis of any one gene requires genome-wide context to set expectations and guide inferences. Here we have used gnomAD data to provide context to aid in the interpretation of human LoF variants.

Targets of approved drugs span a spectrum from highly constrained to completely unconstrained. Why do some genes apparently tolerate pharmacological inhibition but not genetic inactivation? LoF variants, at least in constitutive exons, should affect all tissues for life, whereas drugs differ in tissue distribution and timing and duration of use. Many drugs known or suspected to cause fetal harm are tolerated in adults^43^, and might target developmentally important genes. Constraint is believed to primarily reflect selection against heterozygotes^13^, whose effective gene dosage may differ from that achieved by a drug. Constraint measures natural selection over centuries or millennia; our ancestors’ environment presented different selective pressures than what we face today. Finally, the actions of small molecule drugs do not always map one-to-one onto genes^44–47^. Regardless, these human *in vivo* data show that even a highly deleterious knockout phenotype is compatible with a gene being a viable drug target.

For most genes, the lack of total “knockout” individuals identified to date does not yet provide statistical evidence that this genotype is not tolerated, and, for many genes, such evidence may never be attainable in outbred populations. Bottlenecked populations, individually, are unlikely to yield two-hit individuals for a pre-specified gene of interest, though the sequencing of many different, diverse bottlenecked populations will certainly expand the set of genes accessible by this approach. Identification of two-hit individuals will be most greatly aided by increased investment in the ascertainment and characterization of consanguineous cohorts, where the sample size required for any given gene is often orders of magnitude lower than in outbred populations. Our analysis is limited by sample size, insufficient diversity of sampled populations, and simplifying assumptions about population structure and distribution of LoF variants, so our calculations should be taken as rough, order-of-magnitude estimates. Nonetheless, this strategic roadmap for the identification of human “knockouts” should inform future research investments and rationalize the interpretation of existing data.

Recall-by-genotype efforts to characterize humans with genotypes of interest are only valuable if the variants in question are true LoF. Automated filtering^7^ and transcript expression-aware annotation^14^ are powerful tools, but we demonstrate the continued value of manual curation for excluding further false positives, assessing and interpreting the cumulative allele frequency of true LoF variants, and identifying error modes or biological phenomena that give rise to non-random distributions of pLoF variants across a gene. Such curation is essential prior to any recontact efforts, and indeed, establishing methods for high-throughput functional validation^48^ of LoF variants should be a high priority. Our curation of pLoF variants in neurodegenerative disease genes is limited by a lack of functional validation and detailed phenotyping; in a companion paper we demonstrate a deeper investigation of the effects of LoF variants in *LRRK2*^49^.

As the value of human genetics for drug discovery has been demonstrated repeatedly, we expect that drug development projects will increasingly be accompanied by efforts to study the phenotypes of human carriers of LoF variants. Because the cost of drug discovery is driven overwhelmingly by failure^50^, successful interpretation of LoF data to select the right targets and the right clinical pathways will yield an outsize benefit for research productivity and, ultimately, human health.

## Online Methods

### Data sources

pLoF analyses used the gnomAD dataset of 141,456 individuals^7^. For data consistency, all genome-wide constraint and CAF analyses used only the 125,748 gnomAD exomes. Curated analyses of individual genes used all 141,456 individuals including 15,708 whole genomes. Gene lists used in this study were extracted from public data sources between September 2018 and June 2019. Data sources and criteria for gene list extraction are shown in Extended Data Table 3.

### Calculation of pLoF constraint

The calculation of constraint values for genes has been described in general elsewhere^10,12^ and for this dataset specifically by Karczewski et al^7^. Constraint calculations used LOFTEE-filtered (“high confidence”) single-nucleotide variants (which for pLoF means nonsense and essential splice site mutations) found in gnomAD exomes with minor allele frequency < 0.1%. Only unique canonical transcripts for protein-coding genes were considered, yielding 17,604 genes with available constraint values. For curated genes (Table 2), the number of observed variants passing curation was divided by the expected number of variants to yield a curated constraint value. For *PRNP*, the expected number of variants was adjusted by multiplying by the ratio of the sum of mutation frequencies for all possible pLoF variants in codons 1-144 to the sum of mutation frequencies for all possible pLoF variants in the entire transcript, yielding 6 observed out of 6.06 expected. For *MAPT*, the expected number of variants was taken from Ensembl transcript ENST00000334239, which includes only the exons identified as constitutively brain-expressed in Fig. 3b.

### Calculation of pLoF heterozygote and homozygote/compound heterozygote frequencies

LOFTEE-filtered high-confidence pLoF variants with minor allele frequency <5% in 125,748 gnomAD exomes were used to compute the proportion of individuals without a loss-of-function variant (q); the CAF was computed as p = 1-sqrt(q). This approach conservatively assumes that, if an individual has two different pLoF variants, they are in *cis* to each other and count as only one pLoF allele.

For outbred populations (Fig. 2a), we used the value of p from all 125,748 gnomAD exomes, as this allows the largest possible sample size. This includes some individuals from bottlenecked populations, for which the distribution of p does differ from outbred populations, but these individuals are a small proportion of gnomAD exomes (12.6%). This also includes some consanguineous individuals, but these are an even smaller proportion of gnomAD exomes (2.3%), and any difference in the value of p between consanguineous and outbred populations is expected to be very small. Heterozygote frequency was calculated as 2p(1-p) and homozygote and compound heterozygote frequency was calculated as p^2^. Lines indicate the size of gnomAD (141,456 individuals) and the world populaton (6.69 billion).

For bottlenecked populations (Fig. 2b), we used the value of p from the 10,824 Finnish exomes only. Lines indicate the number of Finns in gnomAD (12,526) and the population of Finland (5.5 million).

For consanguineous individuals (Fig. 2c), we again used the value of p from all gnomAD exomes, because p is not expected to differ greatly in consanguineous versus outbred populations. We used the mean proportion of the genome in runs of autozygosity (a) from individuals self-reporting second cousin or closer parents in East London Genes & Health, a = 0.05766 (rounded to 5.8%). Heterozygote frequency was calculated as 2p(1-p) and homozygote and compound heterozygote frequency was calculated as (1-a)p^2^ + ap. Lines indicate the number of consanguineous South Asian individuals in gnomAD (*N*=2,912, by coincidence the same number as report second cousin or closer parents in ELGH) based on F > 0.05 (a conservative estimate, since second cousin parents are expected to yield F = 0.015625), and the estimated number of individuals in the world with second cousin or closer parents (10.4% of the world population)^9^.

Several caveats apply to our CAF analysis. Our approach naively treats genes with no pLoFs observed as having p=0, even though pLoFs might be discovered at a larger sample size. It also naively treats genes with one pLoF allele observed as having p=1/(2*125748), even though on average singleton variants have a true allele frequency lower than their nominal allele frequency^10^. We naively group all populations together, even though the distribution of populations sampled in gnomAD does not reflect the world population^7^; we believe this is reasonable because CAF for many genes is driven by singletons and other ultra-rare variants for which frequency is not expected to differ appreciably by continental population^10^. It is important to note that the histograms shown in Fig. 2 reflect the expected frequency of heterozygotes and homozygotes/compound heterozygotes, based on gnomAD allele frequency, rather than the actual observed frequency of individuals with these genotypes in gnomAD. Finally, the sample size for all gnomAD exomes (Fig. 2a and 2c) is larger than for only Finnish exomes (Fig. 2b). For a version of Fig. 2 with the global gnomAD population downsampled to the same sample size as the gnomAD Finnish population, see Extended Data Fig. 2.

### Knockout roadmap

For the knockout “roadmap” (Fig. 2d-e) we classified genes according to the current status of human disease association and LoF ascertainment. Genes were classified as having a Mendelian disease association if they were present in OMIM with the filters described in Extended Data Table 1.

Remaining genes were classified as “2-hit LoF reported” based on presence in one or more of the following gene lists: homozygous LoF genotypes in gnomAD curated as described^7^; filtered homozygous LoF genotypes in runs of autozygosity with minor allele frequency <1% in canonical transcripts in the Bradford, Birmingham, and ELGH^25^ cohorts (total *N*=8,925); observed number of imputed homozygotes >1 or number of compound heterozygous carriers where minor allele frequency <2% (for both variants) in DeCODE^28^; homozygous LoF reported in PROMIS^27^; homozygous LoF with minor allele frequency <1% in UK Biobank^29^.

The remainder of genes were sequentially classified as “likely haploinsufficient” if pLI > 0.9 in gnomAD, “pLoF not yet observed” if CAF = 0 in gnomAD, and, finally, “pLoF observed in gnomAD” if CAF > 0 in gnomAD.

### Genetic prevalence estimation

Here, we define “genetic prevalence” for a given gene as the proportion of individuals in the general population at birth who harbor a pathogenic variant in that gene that will cause them to later develop disease. Genetic prevalence has not been well-studied or estimated for most disease genes.

In principle, it should be possible to estimate genetic prevalence simply by examining the allele frequency of reported pathogenic variants in gnomAD. In practice, three considerations usually preclude this approach. First, the present gnomAD sample size of 141,456 exomes and genomes is still too small to permit accurate estimates for very rare diseases. Second, the mean age of gnomAD individuals is ~55, above the age of onset for many rare genetic diseases, and individuals with known Mendelian disease are deliberately excluded, so pathogenic variants will be depleted in this sample relative to the whole birth population. Third and most importantly, a large fraction of reported pathogenic variants lack strong evidence for pathogenicity and are either benign or low penetrance^10,41^, so without careful curation of pathogenicity assertions, summing the frequency of reported pathogenic variants in gnomAD will in most cases vastly overestimate the true genetic prevalence of a disease.

Instead, we searched the literature and very roughly estimated genetic prevalence based on available data. In most cases, we took disease incidence (new cases per year per population), multiplied by proportion of cases due to variants in a gene of interest, multiplied by average age at death in cases. In some cases, estimates of at-risk population or direct measures of genetic prevalence were available. Details of the calculations undertaken for each gene are provided in Extended Data Table 2.

### Data and source code availability

Analyses utilized Python 2.7.10 and R 3.5.1. Data and code sufficient to produce the plots and analyses in this paper are available at https://github.com/ericminikel/drug_target_lof

## Acknowledgments

This study was performed under ethical approval from the Partners Healthcare Institutional Research Board (2013P001339/MGH) and the Broad Institute Office of Research Subjects Protection (ORSP-3862). We thank all of the research participants for contributing their data. EVM acknowledges support from the National Institutes of Health (F31 AI122592) and an anonymous organization. gnomAD data aggregation was supported primarily by the Broad Institute, gnomAD analysis was supported in part by NIDDK U54 DK105566, and development of LOFTEE by NIGMS R01 GM104371. The content is solely the responsibility of the authors and does not necessarily represent the official views of the National Institutes of Health. ELGH is funded by the Wellcome Trust (102627, 210561), the Medical Research Council (M009017), Higher Education Funding Council for England Catalyst, Barts Charity (845/1796), Health Data Research UK (for London substantive site), and research delivery support from the NHS National Institute for Health Research Clinical Research Network (North Thames). NW is supported by a Rosetrees and Stoneygate Imperial College Research Fellowship. The results published here are in part based upon data: 1) generated by The Cancer Genome Atlas managed by the NCI and NHGRI (accession: phs000178.v10.p8). Information about TCGA can be found at http://cancergenome.nih.gov, 2) generated by the Genotype-Tissue Expression Project (GTEx) managed by the NIH Common Fund and NHGRI (accession: phs000424.v7.p2), 3) generated by the Exome Sequencing Project, managed by NHLBI, 4) generated by the Alzheimer’s Disease Sequencing Project (ADSP), managed by the NIA and NHGRI (accession: phs000572.v7.p4). We thank Jaakko Kaprio and Mitja Kurki (Finnish Twins AD cohort) and Academy of Finland grant 312073, and Ruth McPherson (Ottawa Genomics Heart Study) for providing information on individuals with *PRNP* truncating variants. We thank Jeffrey B. Carroll, Karl Heilbron, J. Fah Sathirapongsasuti, and Laurent C. Francioli for comments and suggestions. A subset of the analyses reported here originally appeared as a blog post on CureFFI.org (http://www.cureffi.org/2018/09/12/lof-and-drug-safety/).

## Extended Data

**Extended Data Table 1.** Data sources for gene lists used in this study. For analysis all lists were subsetted to protein-coding genes with unambiguous mapping to current approved gene symbols; numbers in the table reflect this. Note that the gene counts here reflect totals from the full universe of 19,194 genes; some numbers quoted in the main text reflect only the subset of genes with non-missing constraint values.

| List | N | Description |
| --- | --- | --- |
| All | 19,194 | HGNC protein-coding genes<sup>54</sup>. |
| Olfactory receptors | 371 | As reported by Mainland et al<sup>55</sup>. |
| Homozygous LoF tolerant | 330 | Genes with at least two different high-confidence pLoF variants each found in a homozygous state in at least one individual in gnomAD exomes. |
| Autosomal recessive | 527 | OMIM disease genes deemed to follow autosomal recessive inheritance according to extensive manual curation by the Przeworski group<sup>56</sup>. |
| Autosomal dominant | 307 | OMIM disease genes deemed to follow autosomal dominant inheritance according to extensive manual curation by the Przeworski group<sup>56</sup>. |
| Essential in culture | 683 | Genes deemed essential in cultured cell lines based on CRISPR screens<sup>24</sup>. |
| ClinGen haploinsufficient | 294 | Genes with sufficient evidence for dosage pathogenicity (level 3) as determined by the ClinGen Dosage Sensitivity Map<sup>18</sup>. |
| Approved drug targets | 386 | Genes listed as the top-ranked mechanistic target of approved drugs in the DrugBank 5.0 XML release<sup>17</sup> (accessed Sep 12, 2018). Includes products approved by a variety of agencies including FDA, EMA, and Health Canada. Genes were extracted from the XML file using a custom python script with the criteria `target.attrib['position'] == '1'`, `known-action == 'yes'`, and `group == 'approved'`. |
| Positive targets | 143 | Action listed in DrugBank as: activator, agonist, chaperone, cofactor, gene replacement, inducer, partial agonist, positive allosteric modulator, positive modulator, potentiator, or stimulator |
| Negative targets | 243 | Action listed in DrugBank as: antagonist, blocker, degradation, inhibitor, inverse agonist, negative modulator, neutralizer, or suppressor |
| Other & unknown (effect) | 94 | Action not listed in DrugBank, or any action other than those listed above for positive and negative targets. |
| Small molecule | 176 | DrugBank type == 'small' |
| Antibody | 18 | DrugBank type == 'biotech' and DrugBank categories include 'Antibodies' |
| Other (modality) | 35 | DrugBank type == 'biotech' and DrugBank categories do not include 'Antibodies' |
| Oncology | 45 | ATC level 1 code L |
| Cardiovascular | 38 | ATC level 1 code C |
| Endocrine | 24 | ATC level 1 code G or H |
| Metabolic & alimentary | 38 | ATC level 1 code A |
| Neurology | 35 | ATC level 1 code N |
| Respiratory | 12 | ATC level 1 code R |
| Skeletomuscular | 14 | ATC level 1 code M |
| Other (indication) | 29 | ATC level 1 code B, D, J, P, S, or V |
| Rhodopsin-like GPCRs | 689 | HGNC gene set 140: “G protein-coupled receptors, Class A rhodopsin-like”<sup>54</sup>. |
| Ion channels | 326 | HGNC gene set 177: “Ion channels”<sup>54</sup>. |
| Nuclear receptors | 48 | IUPHAR/BPS Guide to Pharmacology “Nuclear receptors” list<sup>57</sup>. |
| Enzymes | 1,178 | IUPHAR/BPS Guide to Pharmacology “Enzymes” list<sup>57</sup>. |
| GWAS hits | 6,336 | Closest gene to GWAS hits with $P < 5\text{-e}8$ in the EBI GWAS catalog (MAPPED_GENE column)<sup>58</sup>. |
| OMIM genes | 3,367 | Associated to a phenotype with a 6-digit MIM number assigned, no qualifying ‘?’, ‘{’ or ‘[’ in the phenotype description, and the words “response”, “susceptibility”, and “somatic” absent from the phenotype description, in the OMIM Synopsis of the Human Gene Map (morbidmap.txt) as of June 11, 2019. |
| All (tissue expression) | 7,931 | Expressed at >1 TPM in all 53 tissues in GTEx v7 |
| Some (tissue expression) | 9,698 | Expressed at >1 TPM in >0 and <53 tissues in GTEx v7 |
| None (tissue expression) | 1,076 | Expressed at >1 TPM in 0 tissues in GTEx v7 |
| Mouse heterozygous lethal knockout | 401 | Human orthologs mapping to at least one of the 404 genes with a lethal heterozygous phenotype reported in at least one knockout mouse line per MouseMine<sup>23</sup>. |

**Extended Data Table 2.** Estimation of genetic prevalence for gain-of-function genetic neurodegenerative diseases. Data sources were identified through PubMed and Google Scholar searches. Genetic prevalence was defined as the proportion of the population at birth carrying a mutation and destined to later develop disease, and estimated as described for each gene.

| Gene | Basis | Estimate |
| --- | --- | --- |
| *HTT* | A reported HD incidence of 0.38 cases per 100,000 per year based on meta-analysis<sup>59</sup> multiplied by an average age at death of ~60 for the most common CAG lengths<sup>60</sup>. Finally, a genetic screen of a general population sample<sup>61</sup> found ≥40 CAG repeat alleles, which are presumed to be fully penetrant, in 3 individuals out of 7,315, for a genetic prevalence of 1 in 2,438. | 1 in 4,386 |
| *HTT* | Prevalence of 13.7 per 100,000 symptomatic plus 81.6 per 100,000 at 25-50% risk in an exhaustive ascertainment study<sup>62</sup>. Assuming there are twice as many individuals at 25% risk as at 50% risk, then on average 33.3% of the 81.6, or 27.1 per 100,000 have the mutation. Thus, 13.7 + 27.1 = 40.8 per 100,000 individuals have an *HTT* CAG expansion. | 1 in 2,451 |
| *HTT* | A genetic screen of a general population sample<sup>61</sup> found $\geq 40$ CAG repeat alleles, which are presumed to be fully penetrant, in 3 individuals out of 7,315 | 1 in 2,438 |
| *LRRK2* | Based on meta-analysis<sup>63</sup>, Parkinson's disease (PD) has an estimated prevalence of 1,903 per 100,000 at age $\geq 80$, meaning the general population's lifetime risk of PD is $\sim 1.9\%$. It is generally stated that about 10% of PD cases are "familial" and the remainder sporadic; in a diverse worldwide case series, *LRRK2* mutations were found in 179/14,253 (1.3%) sporadic cases and 201/5,123 (3.9%) familial cases<sup>64</sup>, implying that *LRRK2* mutations are present in $\sim 1.6\%$ of all PD cases. Thus, *LRRK2* mutations account for a $1.6\% * 1.9\% = \sim 0.030\%$ lifetime risk of PD in the general population†. | 1 in 3,300 |
| *MAPT* | Estimation of the genetic prevalence of *MAPT* gain-of-function mutations is difficult because pathogenic *MAPT* mutations can present with a variety of clinical phenotypes, and common *MAPT* haplotypes are associated with risk for a variety of different neurodegenerative disorders. We were unable to identify any studies of genetic prevalence nor any large case series for any *MAPT*-associated phenotype. As a crude estimate, we considered that frontotemporal dementia has a reported incidence of 2.7-4.1 per 100,000 per year<sup>65</sup> with typical age at death of perhaps 60, and *MAPT* mutations accounting for 5-20% of familial cases, and familial cases accounting for 40% of all cases<sup>66</sup>. Multiplying all these figures results in range of 0.0032% to 0.020% | 1 in 5,000 – 31,000 |
| *PRNP* | We have recently considered the lifetime risk of genetic prion disease in detail<sup>36</sup>. All forms of prion disease (sporadic, genetic, and acquired) appear to be the cause of death of $\sim 1$ in 5,000 people based on either death certificate analysis or division of disease incidence by the overall death rate. $\sim 10\%$ of cases are attributable to *PRNP* variants with evidence for Mendelian segregation (although additional cases harbor lower-penetrance variants). Thus, we expect a genetic prevalence of 1 in 50,000. On the order of $\sim 1$ in 100,000 people in gnomAD and 23andMe harbor high-penetrance *PRNP* variants<sup>36,41</sup>, although as noted above, we expect these datasets to be depleted compared to the population at birth, because prion disease is rapidly fatal and many individuals in these databases are above the typical age of onset. | 1 in 50,000 |
| *SNCA* | As explained above for *LRRK2*, we assumed a 1.9% lifetime risk of Parkinson's disease (PD) in the general population, with 10% of cases being familial. *SNCA* point mutations, duplications, and triplications all appear to be highly penetrant, and in a familial PD case series these accounted for 103/709 = 15% of individuals<sup>67</sup>. Thus, we estimate that *SNCA* mutations account for a $1.9\% * 10\% * 15\% = 0.00028\%$ risk of PD in the general population. | 1 in 360,000 |
| *SOD1* | *SOD1* mutations are believed to account for $\sim 12\%$ to $24\%$ of familial ALS<sup>68,69</sup> and 1% of sporadic ALS<sup>68,70</sup>. One a meta-analysis found that $\sim 4.6\%$ of ALS is familial<sup>71</sup>, although a figure of 10% is also often used<sup>72</sup>. These figures imply that $\sim 1.5 - 3.3\%$ of all ALS is attributable to *SOD1*. The overall incidence of ALS is reported at $\sim 1.6 - 2.2$ per 100,000 per year<sup>73,74</sup>, so the incidence of *SOD1* ALS might be estimated at $\sim 0.024 - 0.073$ per 100,000 per year. Age at death of $\sim 50$ is around average for many *SOD1* mutations<sup>69</sup>, implying a 1.2 – 3.7 per 100,000 population | 1 in 27,000-83,000 |
|  | prevalence of pathogenic SOD1 mutations. |  |

†It is important to consider for a moment how this figure relates to the penetrance of LRRK2 mutations, as LRRK2 variants appear to occupy a spectrum of penetrance<sup>75</sup>. some variants exhibit Mendelian segregation with disease<sup>76,77</sup>, implying high risk; the G2019S variant is estimated to have ~32% penetrance<sup>78</sup>; and other common variants are risk factors with odds ratios of only ~1.2 estimated through genome-wide association studies (GWAS)<sup>79</sup>. The GWAS-implicated common variants were not included in the case series on which our estimate is based<sup>64</sup>, but G2019S does account for the majority of cases in that series. Because the 0.03% estimate here is based on counting symptomatic cases rather than asymptomatic individuals, it will appropriately underestimate the number of G2019S carriers. In essence, in this calculation each G2019S carrier in the population only counts as 1/3 of a person, because they have only a 1/3 probability of developing a disease. It is therefore appropriate that our estimate of genetic prevalence (0.03%) is actually lower than double the allele frequency of G2019S in gnomAD (0.1%).

**Extended Data Table 3.**
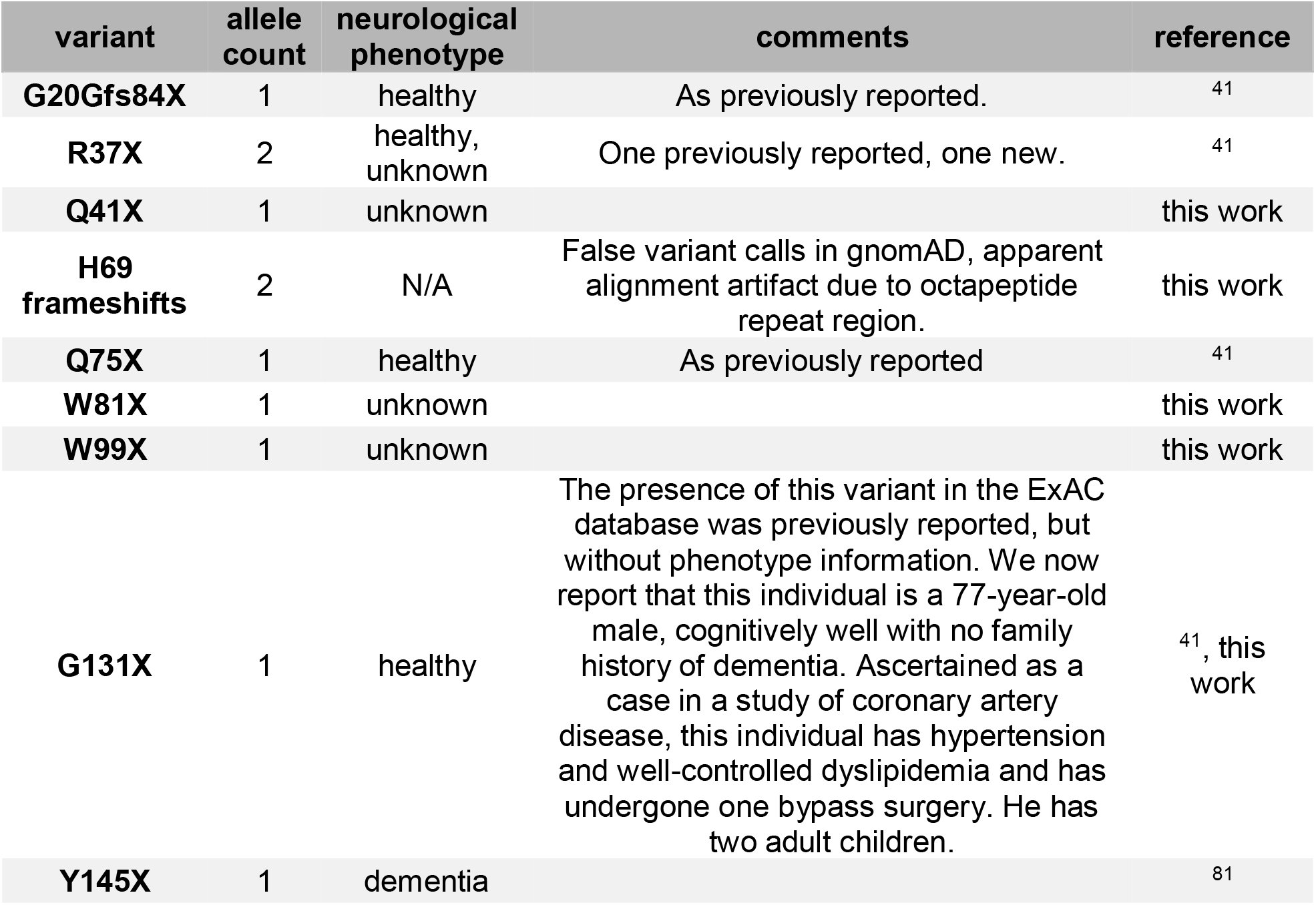
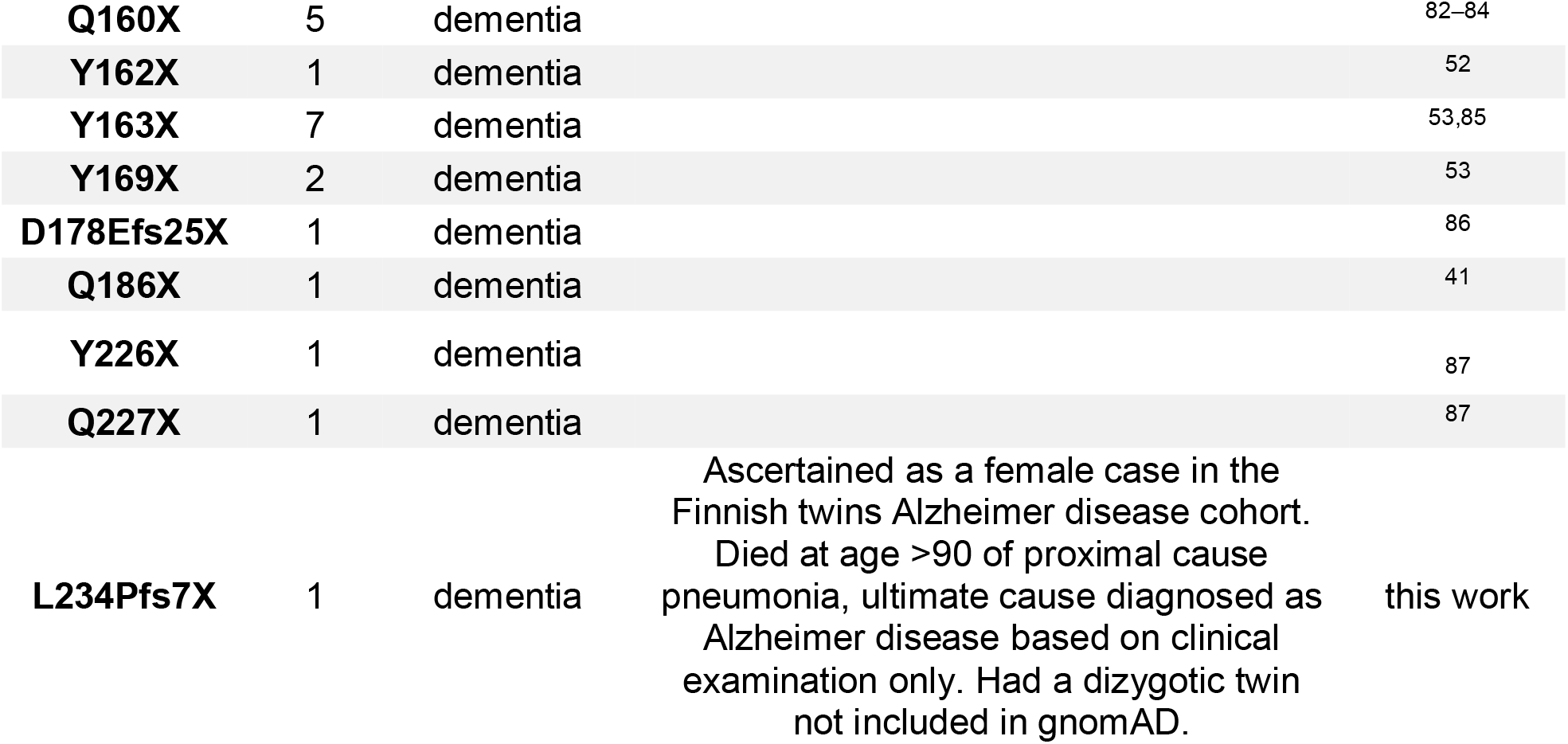
Details of PRNP truncating variants. Allele count for variants from the literature in Fig. 3c is the total number of definite or probable cases with sequencing performed in the studies cited in this table. The L234Pfs7X variant changes PrP’s C-terminal GPI signal from SMVLFSSPPVILLISFLIFLIVGX to SMVPSPLHLX. This novel sequence does not adhere to the known rules of GPI anchor attachmen*t*^80^: GPI signals must contain a 5-10 polar residue spacer followed by 15-20 hydrophobic residues. Thus, this frameshifted PrP would be predicted to be secreted and thus may be pathogenic, explaining the Alzheimer disease diagnosis in this individual. However, it is also possible that the novel C-terminal sequence found here interferes with prion formation, and/or that this variant is incompletely penetrant, and that the diagnosis of Alzheimer’s disease in this individual is merely a coincidence.

**Extended Data Figure 1.**
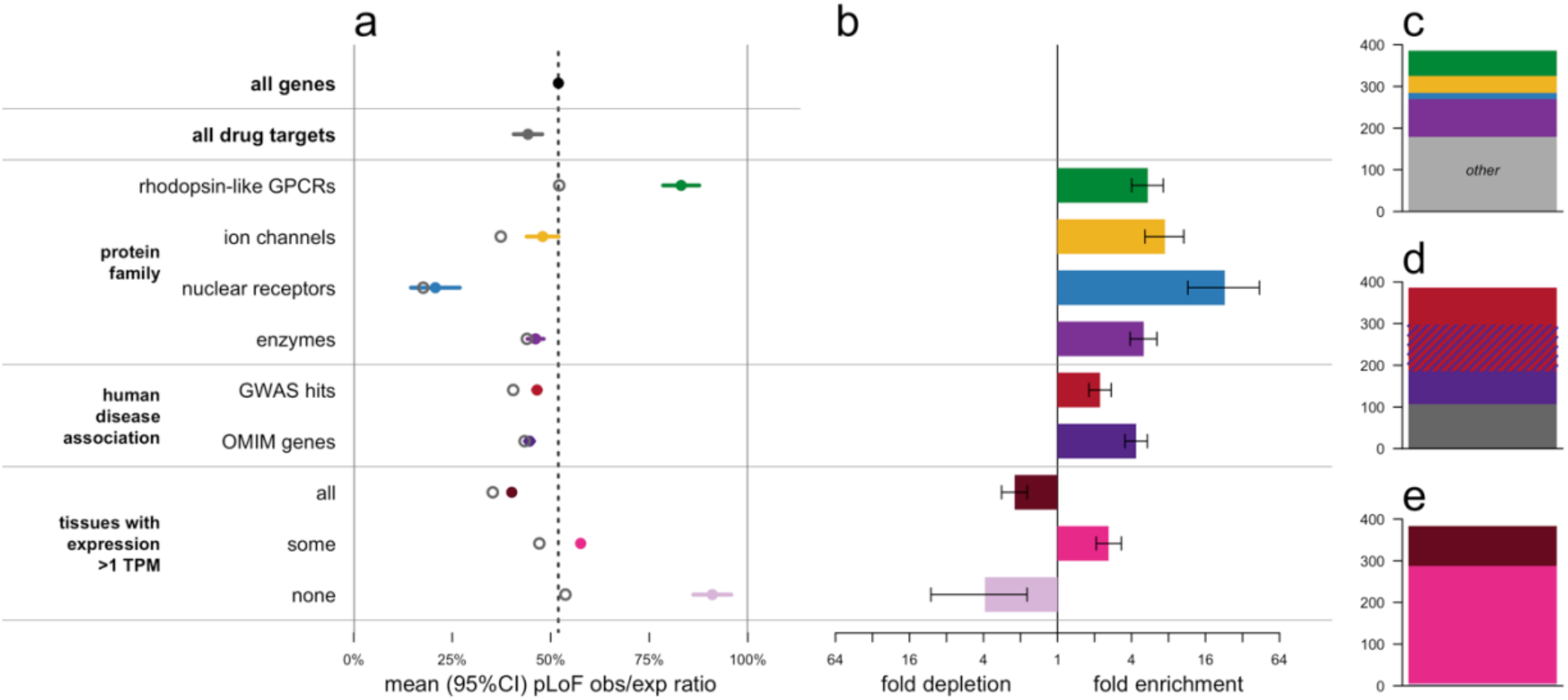
Drug target gene set confounding. a) LoF obs/exp ratios differ significantly from the set of all genes for four canonically “druggable” protein families (top), human disease-associated genes (middle), and genes by broadness of tissue expression (bottom). Within each class, the genes that are drug targets have a lower mean obs/exp ratio (hollow gray circles) than the class overall. b) The “druggable” protein families, disease-associated genes, and genes expressed in some tissues but not others are enriched several-fold among the set of drug targets. c-e) Composition of drug targets when broken down by c) protein family, d) disease association, or e) broadness of tissue expression. The enriched classes account for most drug targets. In a linear model, after controlling for protein family, disease association status, and number of tissues with expression >1 TPM, drug targets are still more constrained than other genes (−8.0% obs/exp, P=0.00012), but the probable existence of additional unobserved confounders cautions against over-interpretation of this observation (see main text).

**Extended Data Figure 2.**
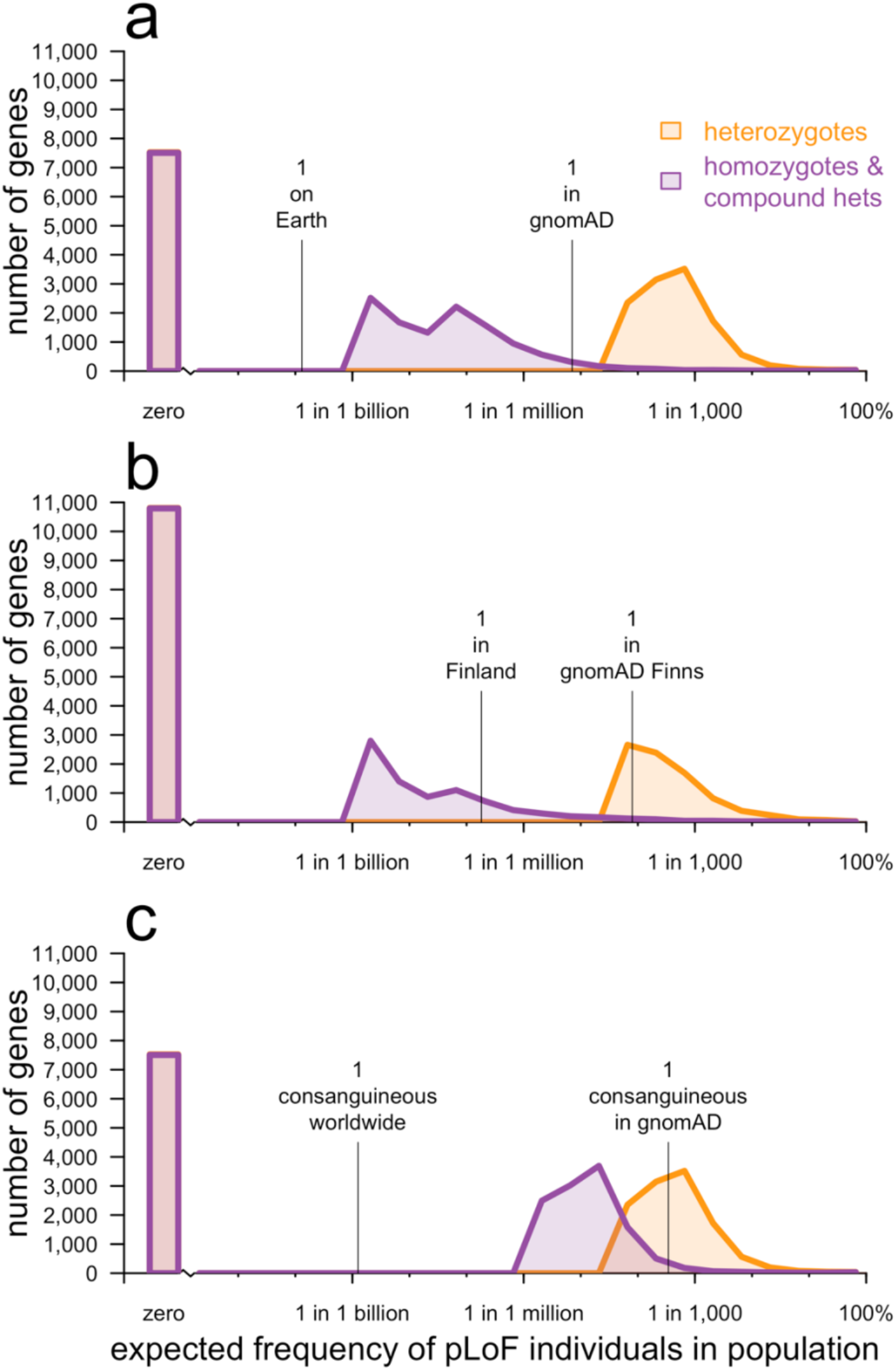
Expected frequency of individuals with one or two null alleles for every protein-coding gene across different population models, with sample size held constant. This is identical to Fig. 2 except as follows. As noted in Online Methods, one caveat about Fig. 2 is that the sample size is larger for the plots using all gnomAD exomes (Fig. 2a and 2c) than for Finnish exomes (Fig. 2b). This figure shows the same analysis, but with the global gnomAD population downsampled to 10,824 randomly chosen exomes so that the same size is identical to that of Finnish exomes. Computation of p = 1-sqrt(q) as described in Methods is computationally expensive for downsampled datasets because it requires individual-level genotypes. Instead, this analysis uses “classic” CAF, which is simply the sum of allele frequencies of all high-confidence pLoF variants each at allele frequency <5%, capped at a total of 100%, for both global and Finnish exomes. The results show that even when sample size is held constant, the number of genes with zero pLoF variants observed is higher in a bottlenecked population than in a mostly outbred population. A constant y axis with no axis breaks is used in this figure to make this difference more clearly visible.

## Supplementary Information

**Supplementary Table 1.**
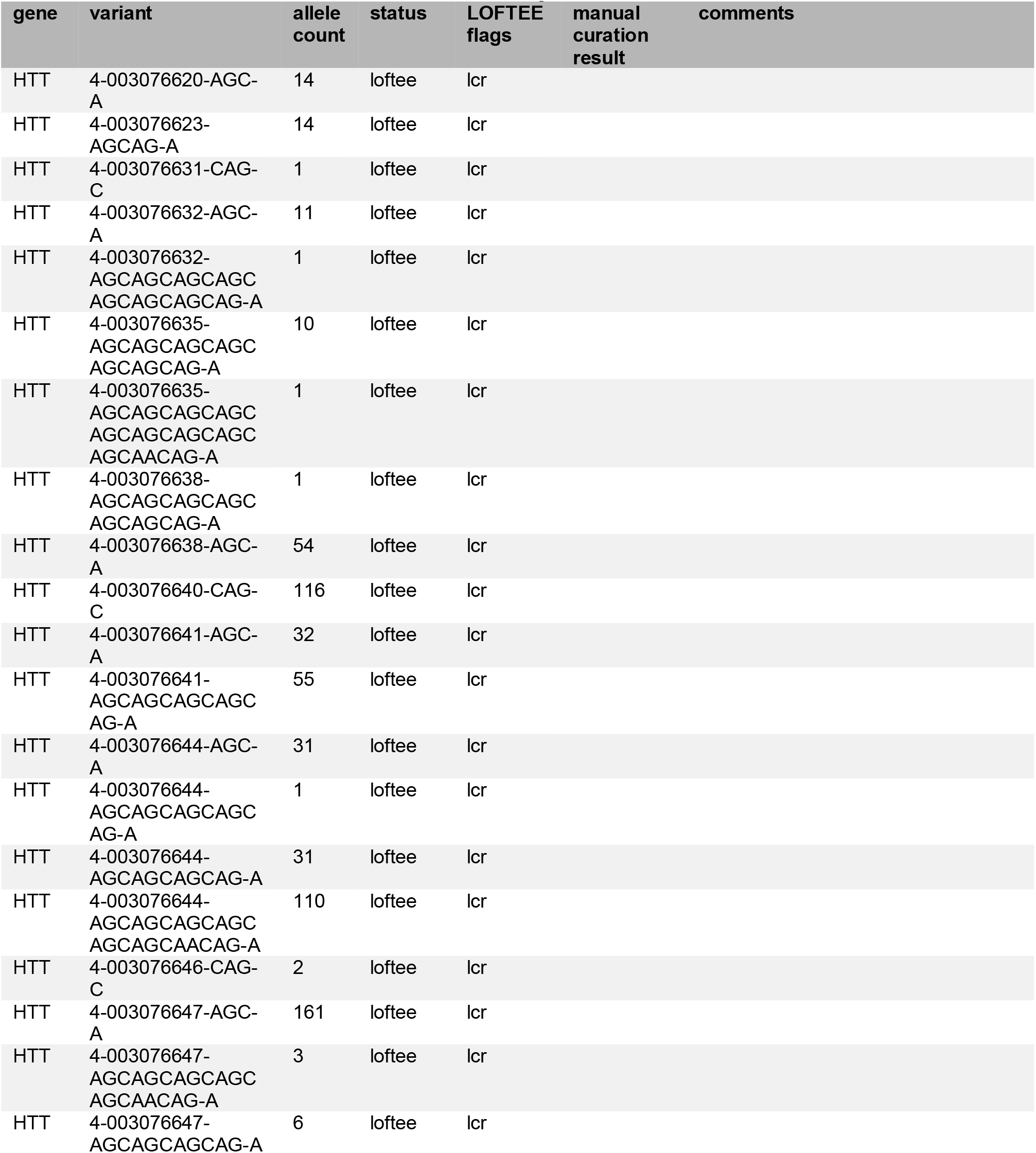
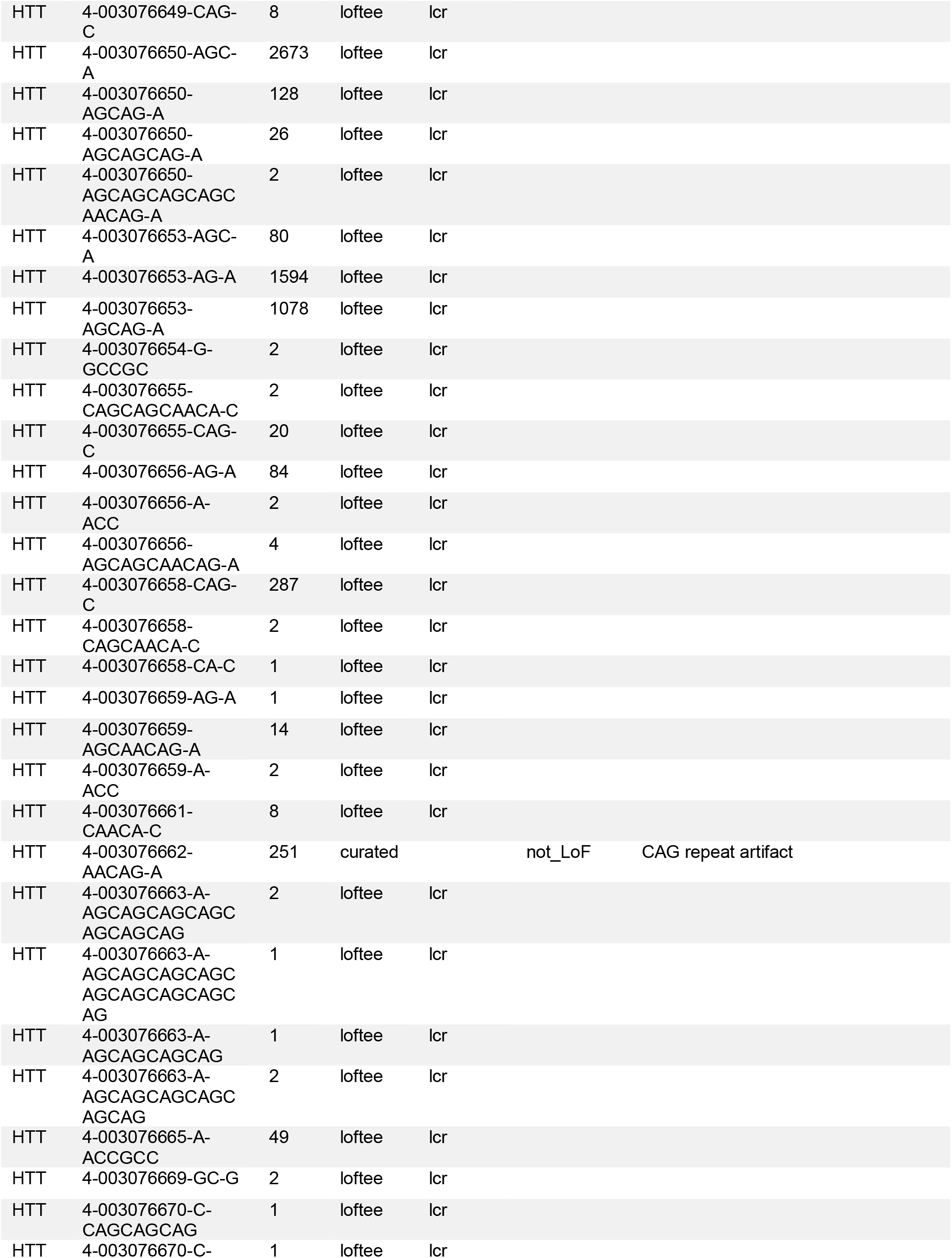
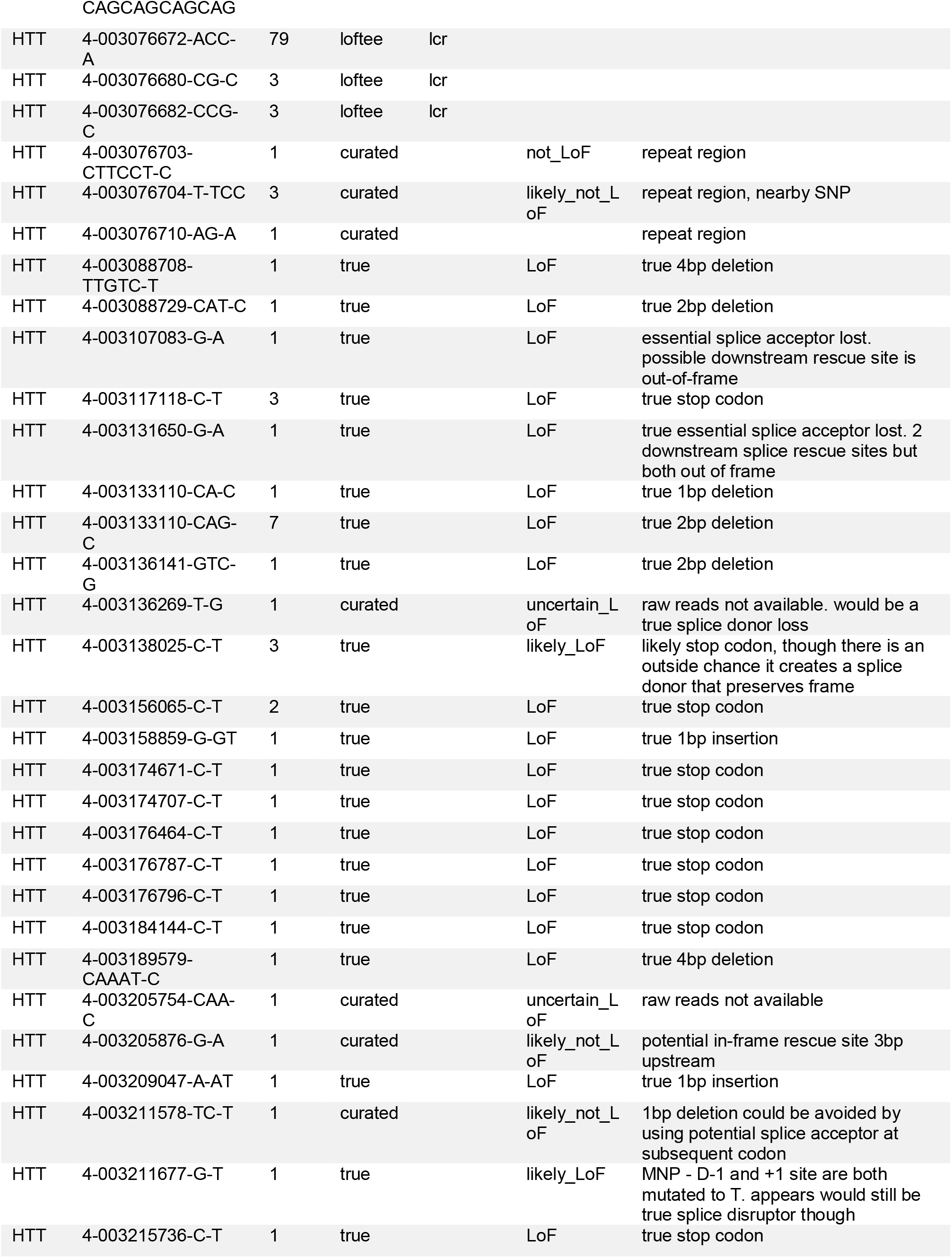
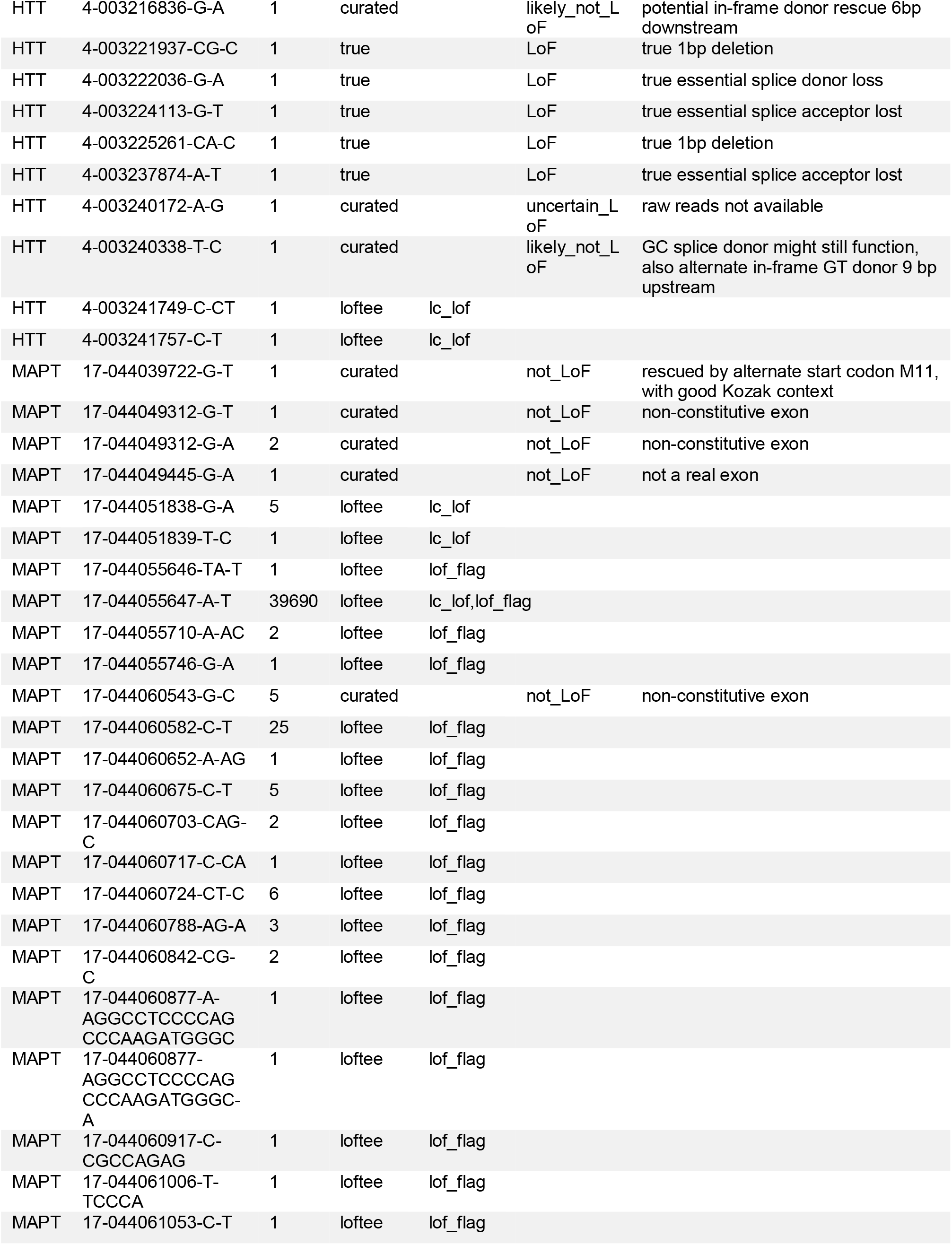
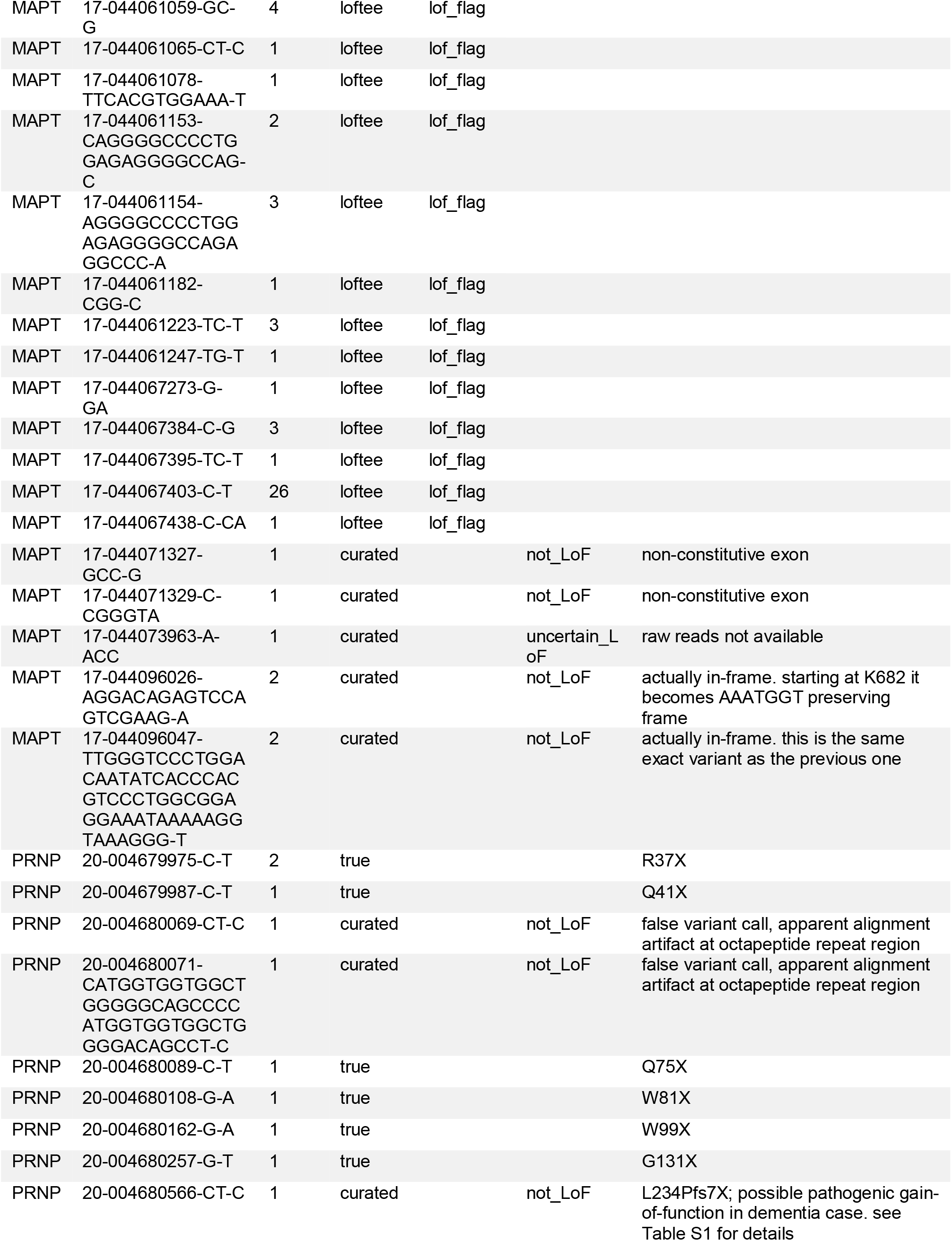
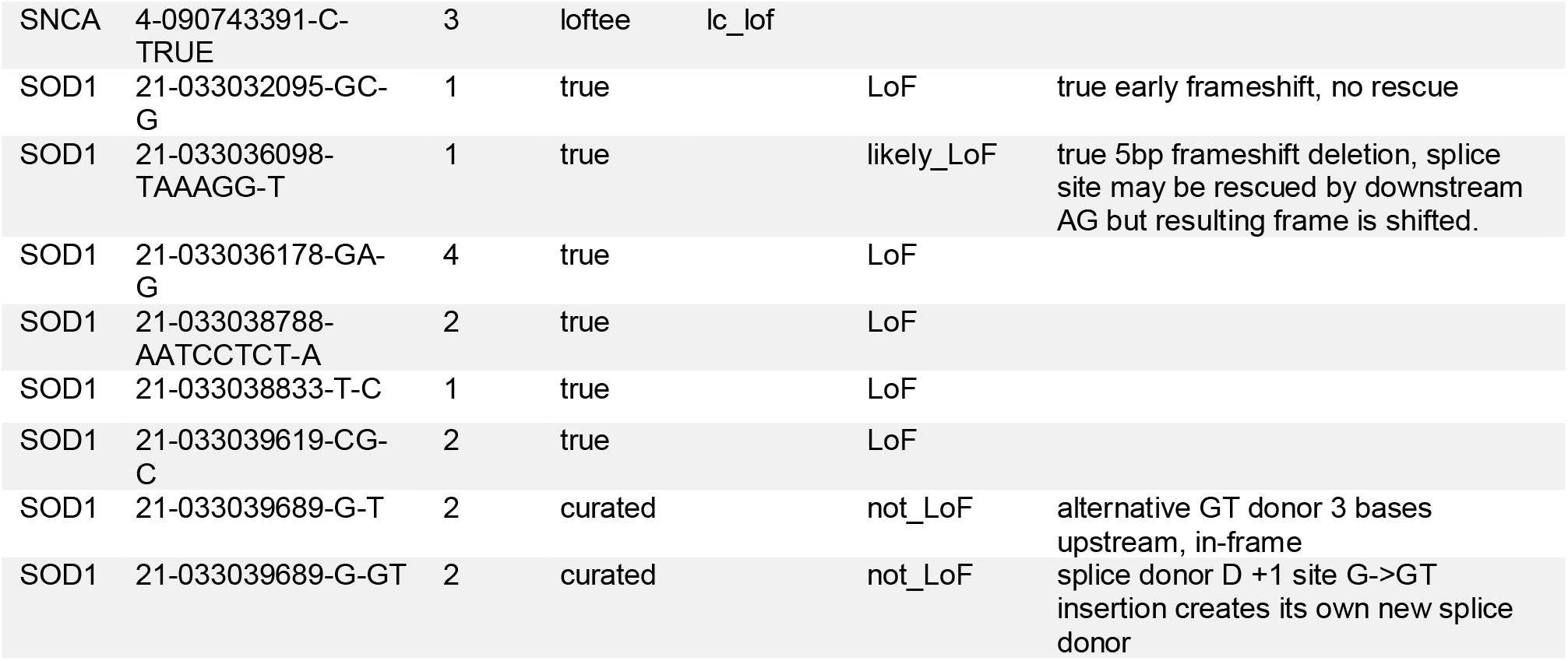
Details of curated variants in neurodegenerative disease genes. LRRK2 is not included here as curation is reported in detail in a separate publicatio*n*^54^. We note that frameshift mutations in SOD1 at codons 126 or 127 have been reported to cause a pathogenic gain-of-function leading to ALS^122,123^. Both of these codons occur in the gene’s fifth and final exon; all of the variants curated as leading to loss-of-function here are in exons 1-4.

